# Thickness and quality controlled fabrication of fluorescence-targeted frozen-hydrated lamellae

**DOI:** 10.1101/2024.07.04.602102

**Authors:** Daan B. Boltje, Radim Skoupy, Clémence Taisne, Wiel H. Evers, Arjen J. Jakobi, Jacob P. Hoogenboom

**Affiliations:** Department of Imaging Physics, Delft University of Technology, Delft, The Netherlands; Kavli Institute of Nanoscience, Delft University of Technology, Delft, The Netherlands; Delmic B.V., Delft, The Netherlands

**Keywords:** Thickness control, Frozen hydrated lamella, Quality control, cryo-ET

## Abstract

Cryogenic focused ion beam (FIB) milling is essential for fabricating thin lamella-shaped samples out of frozen-hydrated cells for high-resolution structure determination. Structural information can only be resolved at high resolution if the lamella thickness is between 100 and 200 nm. While the lamella fabrication workflow has undergone significant improvements since its conception, quantitative, live feedback on lamella thickness and quality is still lacking. Taking advantage of a coincident light microscopy integrated into the FIB-SEM, we present three different strategies that together allow accurate, live control during lamella fabrication. First, we combine 4D-STEM with fluorescence microscope (FM) targeting to determine the lamella thickness. Second, with reflected light microscopy (RLM) we screen target sites for ice contamination and monitor lamella thickness and integrity of the protective Pt coating during FIB milling. Third, we exploit thin-film interference to obtain fine-grained feedback on thickness uniformity below 500 nm. We finally present a full workflow for fluorescence-targeted and quality controlled fabrication of frozen-hydrated lamellae, benchmarked with excellent agreement to energy filtered transmision electron microscopy (EFTEM) measurements and reconstructed tomograms obtained with electron cryo-tomography.

## 1 Motivation

The composition, conformation, and function of most macromolecular complexes depend on their cellular context and must be studied inside cells. Few cells are sufficiently thin to permit direct imaging with cryo-EM. Focused-ion-beam milling enables cryo-EM to visualize macromolecules in cells at high resolution by generating thin sections of frozen hydrated cells. We show how thin cellular sections can be prepared in a controlled fashion using an integrated light microscope coincident with electron and ion beams. The procedure provides live feedback on the thickness and uniformity of the prepared lamella, reducing complexity, and increases the success rate. Combined with its ability for fluorescence-based targeting, our procedure paves the way towards a fully automated workflow for routine fabrication of frozen-hydrated cellular sections.

## 2 Introduction

Cryogenic electron tomography (cryo-ET) has become an integral technique in the quest for a mechanistic understanding of complex, biological processes at the molecular scale, as it remains the sole imaging method capable of discerning the intricate structural features within a cell without labeling [46, 36, 38]. Its utility for gaining biological insights is, however, often hampered by constraints in sample preparation, in particular the need for thin and ideally artifact-free cellular sections. Sufficiently thin sections can be fabricated using a focused ion-beam scanning electron microscope (FIB-SEM), where grazing incident ion bombardment locally removes cellular material to unveil a cross-section of the cell’s interior (lamella), priming it for high-resolution imaging with a transmission electron microscope (TEM) [6, 29, 40, 14]. To resolve the structural information at high resolution, the ideal lamella thickness is limited to around 100 to 200 nm, to remain well below the inelastic mean-free path of electrons in vitreous water ice for TEM imaging at 200 to 300 nm. [41, 44].

In recent years, various improvements and refinements have been made to the cryogenic focused ion beam (cryo-FIB) milling workflow improving throughput, reliability, sample yield and quality. Gas injection systems (GISs) in the cryo-FIB microscope allow deposition of a Pt layer protecting the target region from ion exposure. Integrity of this Pt layer during milling is crucial for obtaining thin lamellae. Different approaches to *in-situ* fluorescence imaging have been integrated into the cryo-FIB workflow for selecting target cells and identification of the region of interest for milling [30, 43, 35, 4, 1, 13, 32, 37, 3, 22, 19]. Precise localization of the target in cryo-FIB coordinates is however still challenging due to registration errors and aberrations that result from a refractive index mismatch during fluorescence microscopy [23]. *In-situ* feedback during cryo-FIB milling on lamella quality, specifically its thickness and uniformity, the state of the protective Pt layer, and the inclusion of the biological target, could provide a significant improvement of fabrication yield in the frozen-hydrated lamella workflow.

The lamella thickness and uniformity can be gauged through the use of the scanning electron microscope (SEM) [39, 7]. However, a prerequisite is that the lamella is composed of a homogeneous material, which for cellular samples, is not the case. Moreover, most methods require independent calibration prior to each imaging session and suffer from practical disadvantages [31]. Quantitative thickness estimations for cellular specimen have been reported when imaging the periphery of a cell in the TEM [20], but have not been extended to the lamella fabrication workflow. Alternatively, using quantitative 4D STEM (q4STEM) imaging we recently showed that the lamella thickness can be robustly estimated directly in the FIB-SEM without additional calibrations if the instrument setup allows transmission imaging [31].

Up to now, routine fabrication of sufficiently thin frozen-hydrated sections remains a challenge due to a lack of direct, quantitative feedback on lamella thickness and its uniformity as well as the state of the protective cryo-deposited Pt GIS layer.

Here, we utilise a coincident FM-FIB-SEM setup [3] for automated fluorescence-targeted lamella preparation and present three complementary techniques to estimate the thickness of a frozen hydrated lamella during the milling process without requiring any prior calibration at the start of the milling session. We (i) apply the previously presented q4STEM method to frozen-hydrated lamella, and benchmark the measured thickness against EFTEM, (ii) image the lamella from the foil side using RLM and obtain a thickness estimation from the known milling geometry, and (iii) exploit thin-film interference on a per-pixel basis yielding a fine-grained thickness map of the lamella, thus providing quantitative feedback on the lateral thickness variations. In addition, we show how to account for axial scaling effects caused by the refractive index mismatch (RIM) when selecting targets in fluorescence microscopy and implement this in our approach for automated milling of fluorescent targets. Finally, we combine all of the above in showcasing a thickness and quality-controlled, fluorescence-targeted lamella fabrication workflow.

## Results

### Fluorescent targeting under refractive index mismatch

In our three-beam coincident setup, cryo-fluorescence microscopy is used to identify targets in frozen hydrated cells and surrounding material is subsequently ablated with the FIB to create a lamella containing the target. Given the large temperature difference between the optical objective (at room temperature) and the sample held at 100 K, a dry objective lens (*n*_1_ = 1.0) is used to image into cellular material, which mostly consists of water (*n*_2_ = 1.28 at *T* = 109 K) [18]. This creates a RIM leading to a depth-dependent deformation in the axial coordinate of the optical microscope [23]. It is crucial to consider this effect when fabricating lamellae around fluorescent targets embedded in frozen-hydrated cells.

To compensate for the axial compression due to the RIM the z-position of the vacuum-specimen interface needs to be accurately known. In our coincident three-beam microscope, the sample holder is oriented such that an incident angle of 10*^◦^* is maintained with the FIB, and consequently the incident angle of the SEM is 118*^◦^*. The optical objective is positioned below the specimen, and images the region of interest (ROI) at right angles through the electron microscope (EM) grid as shown in Figure 1a.

**Figure 1:**
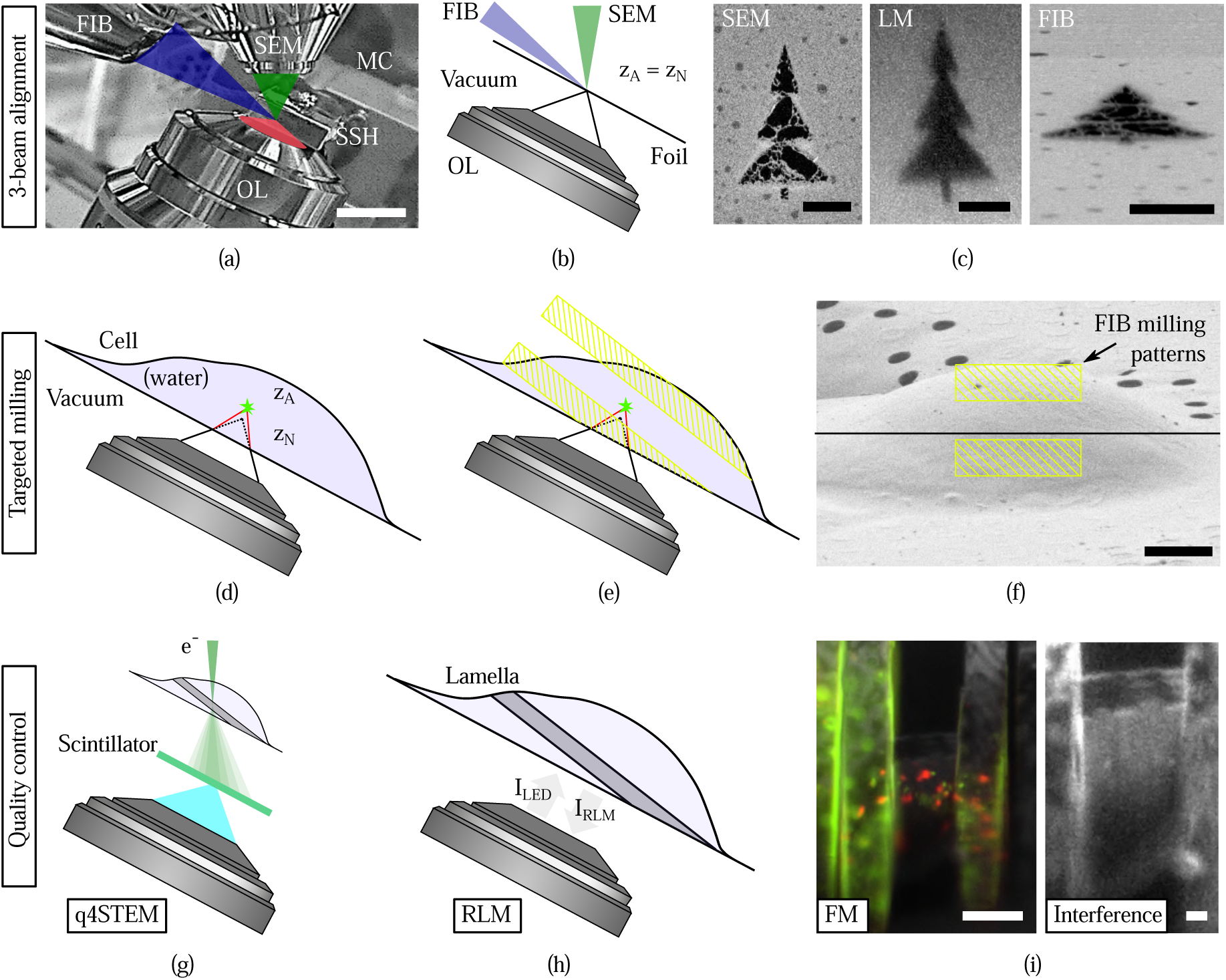
The alignment of the experimental setup used for lamella fabrication (a through c), the targeting & milling approach (d through f) and techniques for quality control (g through i). **(a)** Infrared photograph acquired with the camera mounted on the FIB-SEM host system showing the three coincident beams (SEM, LM and FIB) along with the microcooler and objective lens (OL) mounted inside the microscope chamber. The EM grid containing frozen cells is contained in the sample shuttle holder (SSH) and epi-fluorescence imaging is done from the bottom (transparent red). **(b)** Alignment of the optical microscope is done by moving the objective lens, without a RIM present. The AutoGrid and coverslip have been omitted for clarity. **(c)** Milled pattern imaged with three imaging modalities after alignment, the light microscopy image is acquired by collecting reflected light from the sample. **(d)** When targeting fluorescence inside a cell, the axial coordinate is distorted due to RIM. The measured axial position (*z_N_*) of the fluorescent emitter does not coincide with the actual position (*z_A_*). **(e)** The bottom milling pattern is positioned close to the coincident point, as the fluorescent emitter *always* sits further away from the objective lens due to axial scaling. **(f)** Placement of the asymmetric milling pattern with respect to coincidence (black line), in the FIB field of view prior to rough milling a cellular structure. **(g)** Schematic diagram showing the geometry and detection principle used to measure lamella thickness with q4STEM [31]. By collecting scintillation light from the scattered electron pattern with the optical system, the lamella thickness can be determined. **(h)** Schematic showing the lamella orientation with respect to the objective lens. The lamella thickness can be estimated by collecting the reflected light intensity from the lamella (*I_RLM_*), both through the known imaging geometry and interference effects, see also **(i)**, right. **(i)** The fluorescence intensity can be monitored during and in between milling, thus the presence of the biological target inside the lamella is guaranteed. Thin film interference effects are present in RLM, as seen by the intensity gradient going from bottom (dark) to the top (bright) of the lamella. In addition, interference fringes are visible originating from a Newton interferometer geometry. Both of these effects can be exploited to determine the thickness of the lamella. Scale bars: **(a)** 2 cm, **(c)** 5 µm, **(f)** 5 µm, **(i)** 10 µm (left), 2 µm (right).

With an AutoGrid-mounted sample loaded into the system, the correct height for three-beam coincidence is initially achieved by moving the objective lens to the coincident point of the FIB-SEM. To determine the z position of the air-specimen interface, we use the objective stage to focus onto the grid foil while carefully avoiding ice in the optical path to prevent a RIM. This procedure ensures that the measured axial (microscope or nominal, *z_N_*) position of the grid foil corresponds with the actual position *z_A_* (Figure 1b). This yields Figure 1c, where an alignment pattern is imaged with SEM, FIB and light microscope (LM). The light microscopy image is acquired by collecting reflected light from the sample.

Given the numerical aperture (NA) of 0.85 and the aforementioned refractive indices of objective lens and sample, the axial scaling behaves linearly when fluorescent emitters reside no deeper than ∼9 µm [23]. The upper thickness limit for obtaining vitreous, plunge-frozen hydrated cells, is 10 µm [42], and the acquired focal shift (difference between the actual and measured emitter position) for a targeted emitter position hence ranges from 0 to ∼2.7 µm, scaled linearly at ∼300 nm shift per micrometer emitter depth. This is depicted schematically in Figure 1d, where a fluorescent object at actual depth *z_A_* is aberrantly imaged at the axial coordinate *z_N_* by the optical microscope [23].

Quantitative correction of the focus shift requires precise measurement of the scaled distance, which is possible by determining the focal length difference between the foil (RIM interface) and fluorescent emitter. As lamella fabrication takes place in a step-wise process, with lamella thickness after rough milling ranging around 2 to 3 µm, we opted for a different approach. When imaging from low (*n*_1_ = 1) to high (*n*_2_ = 1.28) refractive index, the axial coordinate system of the optical microscope appears compressed. This allows for an asymmetric milling pattern when the rough cut is made, by placing the bottom pattern a few hundred nanometer away from the coincident point (Figure 1e and f). Consequently, the top milling pattern is placed ∼2.7 µm away from the coincident point, and thus the fluorescent ROI is captured inside the rough cut lamella.

Quality and thickness control is assured through three complementary techniques to estimate the thickness of a frozen hydrated lamella during the milling process. Our previously presented q4STEM method [31] is applied to frozen-hydrated lamella (Figure 1g). Through the known imaging geometry and by exploiting interference effects present in the RLM, the lamella thickness can be estimated too, while also verifying the presence of the fluorescent target inside the lamella using FM (see Figure 1h,i). This will be discussed in more detail below.

### Automated milling of fluorescent targets

We set out to utilise fluorescence targeting to set up automated lamella preparation on selected cellular targets. Our workflow is illustrated in Figure 2, where potential target cells are identified using a low magnification SEM image (white markers) prior to applying the Pt coating with the GIS. Each site is imaged in the FM to refine target positions, and in addition, the sites are also screened in RLM. During the sample preparation, ice crystallites occasionally form, and bigger clusters can obscure the target site. These are often found below the foil side of the grid and thus invisible in the SEM image. They obscure fluorescence imaging and induce additional axial scaling when present directly below the target, and may restrict subsequent tilt series acquisition in the TEM if found along the milling direction of the lamella. RLM imaging using the integrated optical microscope allows identifying and discarding these sites (Figure 2, red marker with white cross), thereby increasing the success rate of high-quality lamella for cryo-ET.

**Figure 2:**
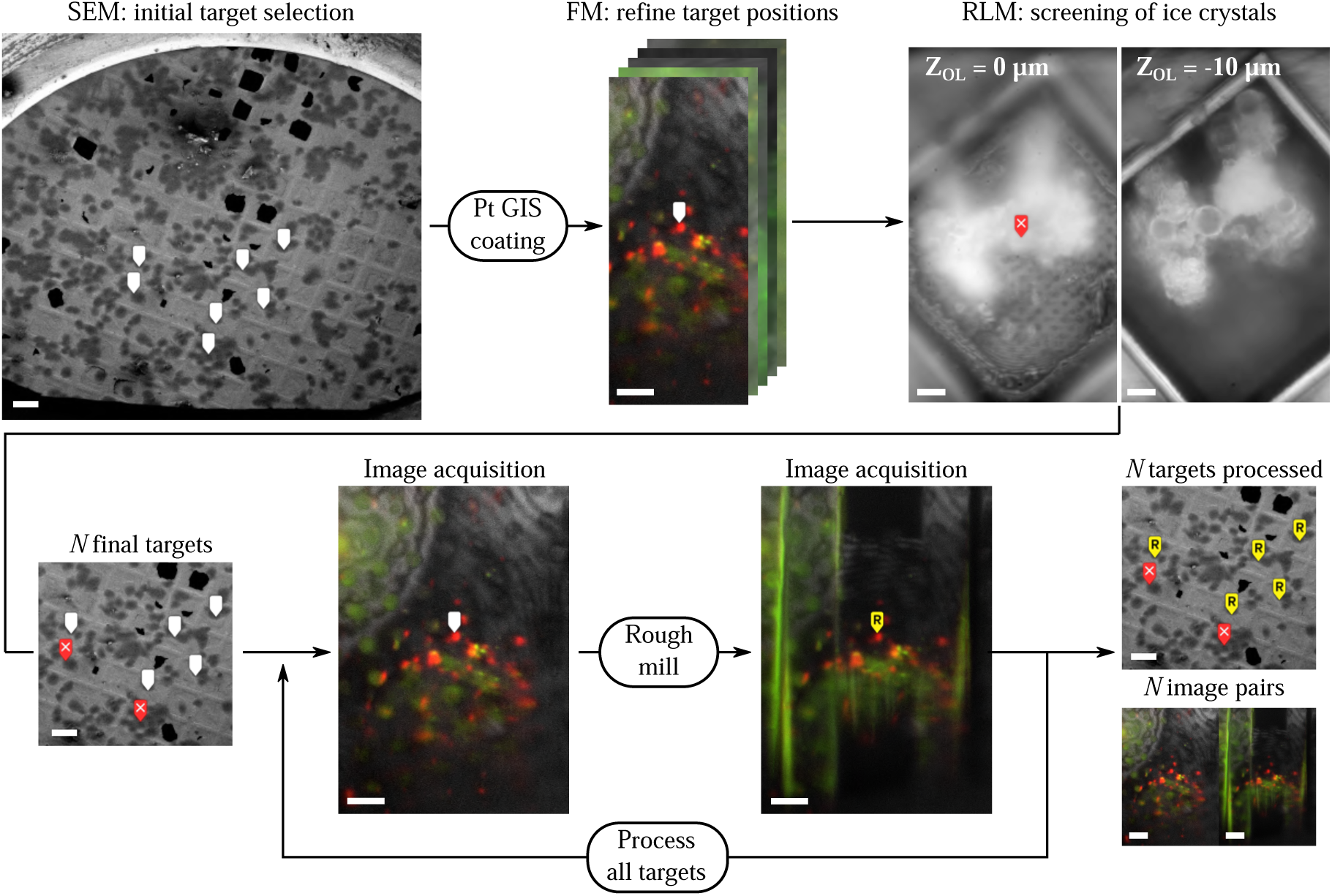
Fluorescence targeted, automated lamella preparation workflow for selected cellular targets. Initial target cells are identified in a low magnification SEM image (white markers, 1 keV, 25 pA, ∼ 2 mm horizontal field width). After applying the Pt coating, the target positions are refined through fluorescence imaging and targets are screened for the presence of ice crystals using the RLM. *N* final targets are rough milled automatically, whilst acquiring FM images before and after. Scale bars: 100 µm (SEM images), 10 µm (RLM images) and 100 µm (FM images).

Once *N* final milling targets have been selected, asymmetric rough milling of the lamella is done in an automated fashion based on the stored *XY Z* stage coordinates obtained from the FM imaging. For each milling site, the Odemis acquisition software positions the stage and automatically acquires images in the relevant channels of the light microscope [28]. This is done prior, and after rough milling, which finally yields *N* milled sites (yellow marker with an R in Figure 2) and image pairs. More examples of these image pairs are shown in Figures S2 and S3, and were acquired over the course of approximately four weeks when milling different biological specimens.

On our Helios Nanolab 650 FIB-SEM (Thermo Fisher Scientific) used in this study, instructions to mill a pre-defined pattern using the FIB, are sent through the XTLib interface (ThermoFisher Scientific) by changing the SEM scan rotation. An iFast script (ThermoFisher Scientific) executes the milling process, after which the scan rotation is set back to the original value, signalling that milling for this site is complete. Similar strategies for controlling beam, stage, and imaging operations can be employed through the instrument API of other manufacturers. The time required to cut stress relief cuts, mill an initial lamella and acquire the images in the LM was about 4 to 5 min per site. The rough lamella can be thinned down to *<*200 nm, whilst having quantitative feedback on the thickness, as outlined below.

### Thickness determination through q4STEM

Reproducible FIB micromachining of high-quality frozen-hydrated lamella requires robust control over the sample thickness during the milling process. Scattering patterns of transmitted electrons can be recorded by turning our integrated light microscope into an optical 2D-STEM detector. As we have shown in previous work, we can achieve this by replacing the ITO-coated glass coverslip of the integrated optical microscope by a scintillator right below the AutoGrid (Figure 3a [3, 31]). While we have used a Yttrium aluminium garnet (YAG) scintillator in our previous proof-of-principle work, its optical emission spectrum in the visible range conflicts with absorption and emission spectra of typical fluorescent targets. We therefore opted for lutetium aluminium perovskite (LuAP) because of its relatively low optical emission wavelength and the fact that it interferes only with 440 nm-centered emission. For recording of electron scattering patterns, the OL is moved down along the SEM optical axis to focus onto the top scintillator surface whilst retaining the alignment with the SEM (Figure 3a). When the estimated lamella thickness is below approximately, 1 µm the thickness can be determined using q4STEM [31]. With the SEM in spot mode, the optical microscope focus is refined using direct electron beam exposure through empty holes in the grid foil (Figure 3b, inset). Individual scattering profiles on different parts of the lamella are then recorded using the SEM beam shift. In our case, temporal coordination of electron exposure is achieved through controlling the Fast Beam Blanker on our Helios Nanolab 650 (Thermo Fisher Scientific) via the transistor-transistor-logic (TTL) signal output from our optical camera. The lamella thickness at each probe position can be determined by radially averaging the scattering profile (Figure 3c) and computing the integrated dark field/bright field (DF/BF) ratio. The DF/BF ratio together with an estimate of the most common scattering angle can then be compared to tabulated data from Monte Carlo (MC) electron scattering simulations to determine a final thickness [31].

**Figure 3:**
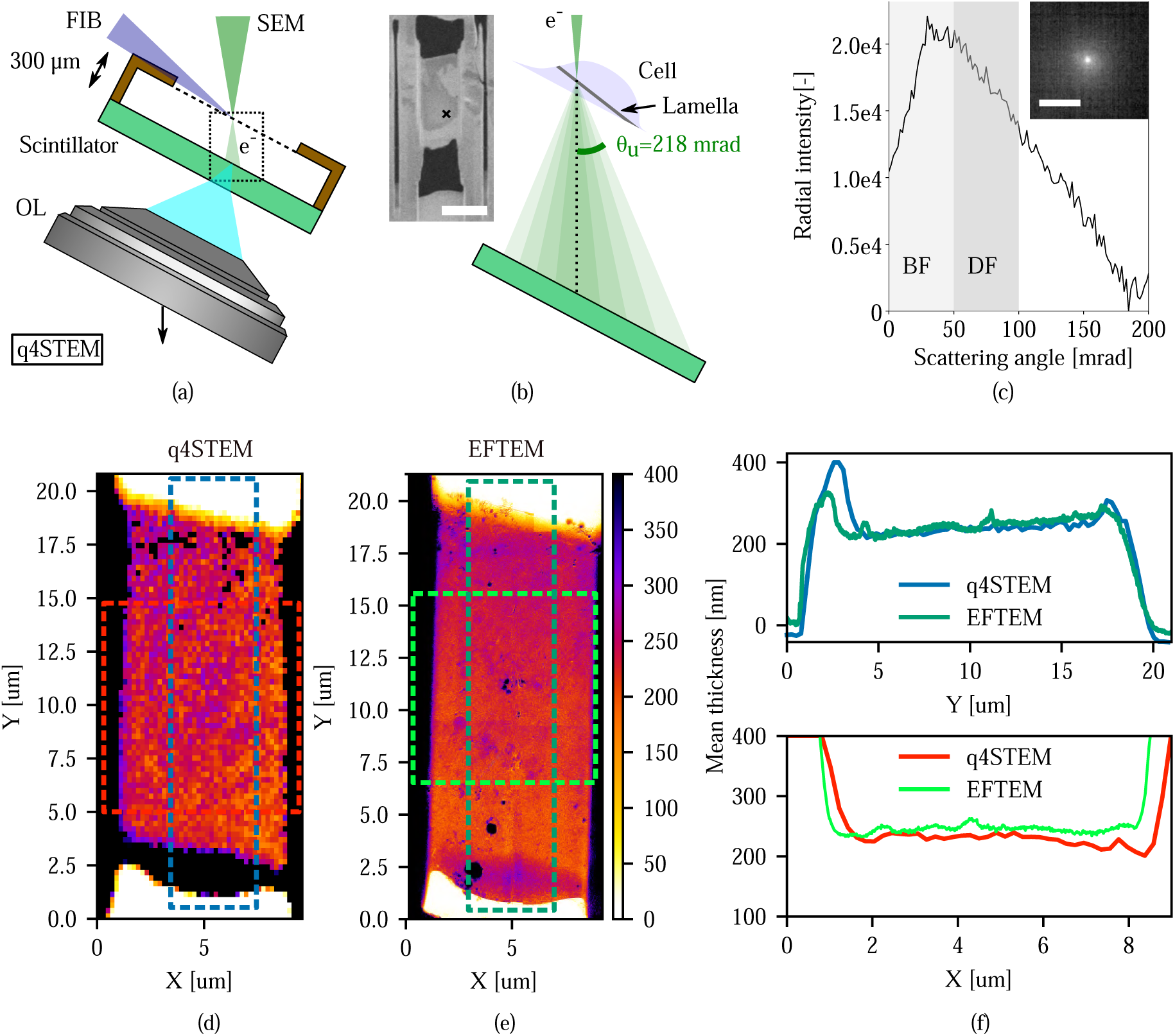
q4STEM-measured thickness of frozen hydrated lamella compared quantitatively with thickness estimates from energy filtered TEM. **(a)** Schematic diagram showing the geometry and detection principle used to measure lamella thickness with q4STEM. The OL focus is moved downward (arrow, along the electron optical axis) to the top scintillator surface, where the scattered electron pattern is visualized by collecting the scintillation light (transparent blue). **(b)** Schematic illustration showing the scattering process in more detail (dotted box in (a)). The left inset shows a SEM image from an intermediate milling step; the black cross marks the position at which the q4STEM measurement was acquired. **(c)** The integrated radial intensity profile as computed from the acquired image (see inset). The thickness is determined by taking the ratio of the virtual dark- and bright field intensity sum, as discussed in earlier work [31]. **(d)** q4STEM thickness map after correcting for the SEM angle of incidence of 52*^◦^*. The map consists of a 64 *×* 100 pixel grid acquired with a 200 nm pixel size. **(e)** EFTEM thickness map from zero-loss imaging after correcting the lamella pre-tilt of 10*^◦^*; pixel size is 3.59 nm. A baseline correction has been applied to the EFTEM data to yield zero thickness outside the lamella (vacuum), **(f)** Mean line profiles along the *X* and *Y* direction of the lamella. The dotted boxes in **(d, e)** annotate the region across which the mean intensity was computed. Scale bars: **(b)** 10 µm, **(c)** 33 µm or 109 mrad.

To validate the ability to robustly estimate thickness from q4STEM measurements on frozen-hydrated lamella, we used fluorescence-guided FIB milling to fabricate a lamella from vitrified HeLa cells stably expressing mRFP-EGFP tandem fluorescent-tagged Microtubule-associated protein 1A/1B light chain 3B (mRFP-GFP-LC3), a dual fluorophore probe for autophagosomes and autophagosome trafficking [17] (Figure S4). Following automatic milling to approximately 250 nm, we acquired a 2D convergent beam electron diffraction pattern at every pixel position of a 2D STEM raster covering the lamella. We then evaluated the resulting 4D STEM dataset using the q4STEM method, resulting in a local thickness at each probe position, corrected for the non-perpendicular incidence of the electron beam (Figure 3d). After transferring the sample to a JEOL JEM3200-FSC TEM operated at 300 kV, we acquired a baseline-corrected EFTEM thickness map using zero-loss imaging (Figure 3e). The two techniques show good agreement in the measured thickness, with a mean difference of 7 nm over the center 4 µm area of the lamella (Figure 3d, e).

Figure 3f shows the mean line profiles in correspondence with the overlays in the respective 2D maps. Both q4STEM and EFTEM profiles display the same ∼50 nm decrease from the Pt-coated leading edge towards the trailing edge of the lamella, caused by the reduced incident ion flux along the length of the lamella due to the grazing incidence. The bump marking an apparent increase in observed thickness at *Y* = 3 µm of the lamella is caused by the platinum-rich GIS coating protecting the lamella during FIB milling, as the scattering profile is assessed using simulated data of pure frozen water.

We assessed whether the low-voltage electron exposure in the SEM led to noticeable beam-induced damage by analyzing a sample of lysosome crystals as described in the Supplemental Information (Figure S5). Our analysis is limited to low order reflections due to the limited sensitivity of the scintillation-based detection method. While a detailed analysis of radiolytic damage extending to higher resolution features will require additional studies with a more sensitive pixelated direct detection STEM detector, we argue thickness determination through q4STEM is a viable route for frozen-hydrated biological samples as: (i) A measurable change in the intensity is only seen for exposures of 100 ms and higher, and q4STEM measurements are performed at lower exposure times. (ii) This diffraction experiment shows that, given the vibrations of the sample holder, it is difficult to expose the same area repetitively. (iii) We apply q4STEM *only* at the edges of the lamella, 2 to 10 µm away from the tomography ROI, and hence no damage to relevant parts is induced. Thus, q4STEM allows precise, local thickness determination on the milled lamella.

### Thickness & quality control using RLM

Live feedback on lamella thickness and quality during milling can be obtained by acquiring RLM images with the coincident light microscope in addition to FM targeting (Figure 4). We illustrate this procedure using fluorescence-targeted lamella preparation from HeLa cells expressing mRFP-GFP-LC3 to simultaneously localise authophagosomes and autophagolysosomes. First, we select ROIs using the fluorescence images (Figure 4a) of suitable cells initially selected using the low magnification SEM image (Figure 4a, inset). The overlay of the two-channel FM image with the RLM simultaneously acquired clearly embeds the fluorescence in the context of the frozen hydrated cell and grid support (Figure 4a). FIB milling is then conducted under a 10*^◦^* angle with the plane perpendicular to the light microscope optical axis leading to the geometry shown in Figure 4b. The optical focus depth is about 500 nm (red shaded area) which makes that we can either focus on the top (Pt, blue), or bottom (foil, green) side of the lamella by moving the sample stage *z*. An estimate of the lamella thickness can then be readily obtained by measuring the width *w* of the bottom (foil) side of the lamella (see inset in (Figure 4b). The lamella thickness then follows from *d* = *w ·* sin(*θ*), where *θ* denotes the milling angle.

**Figure 4:**
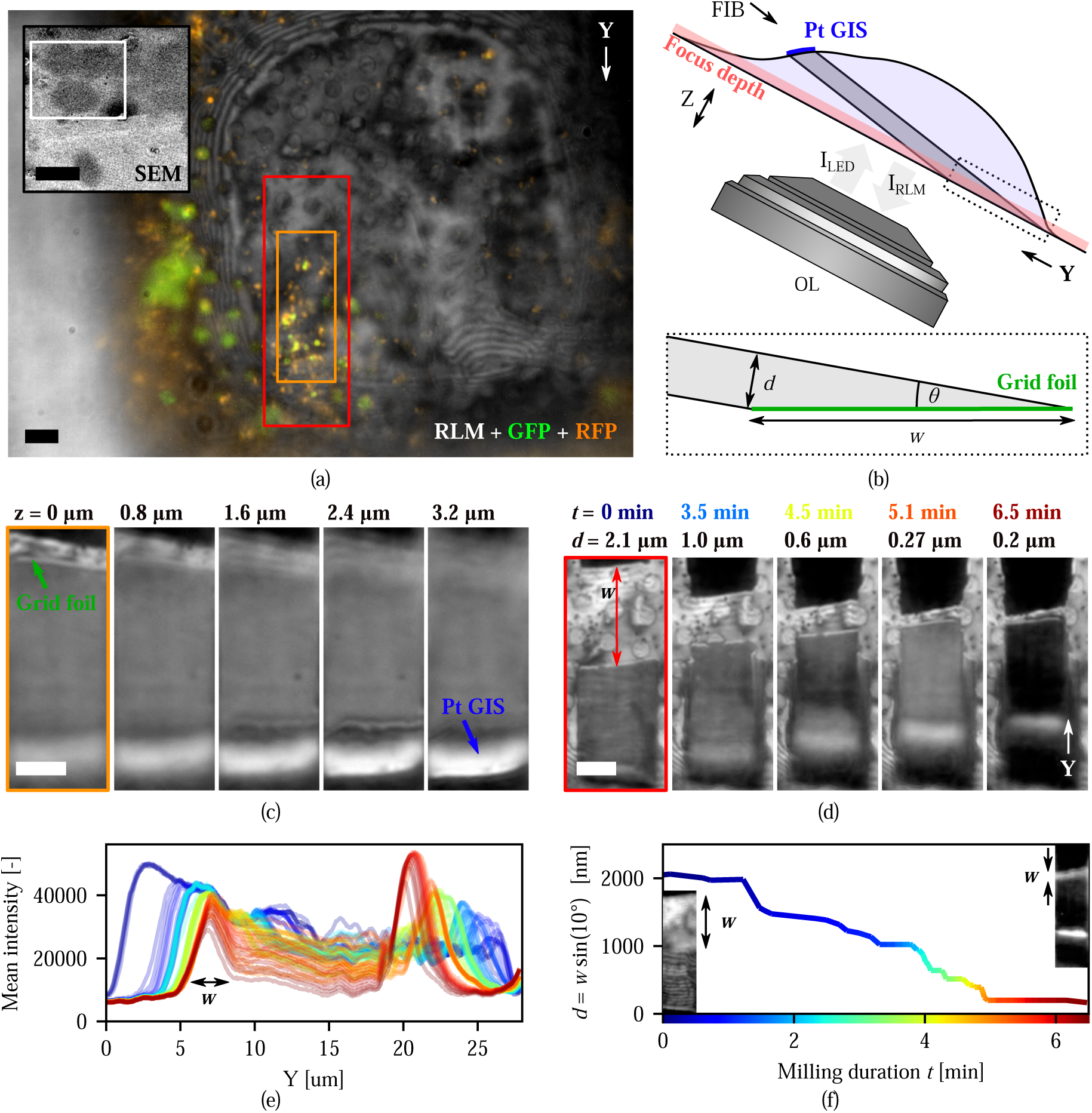
Targeted and quality controlled FIB milling with in-situ fluorescence and reflected light imaging. **(a)** Initial target selection in HeLa cells expressing mRFP-GFP-LC3 using fluorescence microscopy (colors) overlaid on reflected light (grayscale). Green & yellow: autophagosomes and red: autophagolysosomes. The light microscope FOV is denoted by the white rectangle in the top left inset, showing the grid square in the SEM overview image. **(b)** Sketch showing the lamella- and RLM imaging geometry. The optical focus depth (red translucent are) is about 500 nm. The dotted box shows the lamella geometry close to the grid foil. The lamella thickness *d* can be estimated by measuring distance *w* along the tapered wedge of the lamella by *d* = *w ·* sin(*θ*). **(c)** Cropped slices (orange rectangle in (a)) from a RLM z-stack acquired with the lamella milled to about 300 nm in thickness. Top and bottom side of the lamella can be imaged by changing the optical focus (respectively Pt GIS and foil sides, green and blue annotations, also visible in (b)). **(d)** Various stages (time points) of the lamella milling process (FOV cut to red marked area from (a)). The lamella thickness *d* can be estimated by measuring *w* along the bottom wedge of the lamella. **(e)** Mean *X* RLM intensity profiles along the lamella length *Y* (line color indicates milling progression/time according to the color scale in (f)). *w* can be measured from the width of the foil-side intensity peak. **(f)** Estimated lamella thickness *d* versus milling duration. The two insets show the lamella as imaged in RLM at the start and end points of thinning. The lamella thickness can be estimated to approximately 250 nm. Scale bars: **(a)** 5 µm, inset: 50 µm, **(c)** 5 µm, **(d)** 5 µm.

Acquisition of a RLM *z*-stack from the lamella at intermediate milling steps allows quality inspection of the milling process beyond thickness evaluation. As both the foil and Pt GIS side can be brought into focus (Figure 4c), we can with one stack evaluate the (i) lamella thickness *d* by estimating the width *w* from the foil side; (ii) state and uniformity of the Pt GIS layer, protecting the lamella during ion beam bombardment; (iii) uniformity of the milled lamella, and (iv) verification of the milling angle by measuring the defocus (4 µm) along total length (22.4 µm) of the lamella: *θ* = arcsin (4 µm*/*22.4 µm) = 10.2*^◦^*. By setting the focus in between the top and bottom side of the lamella (1.6 µm in Figure 4c), it is possible to both obtain a rough estimate for the lamella thickness *d*, while the highly reflective contribution from the Pt GIS layer can be used to assess the uniformity of the protective layer.

Snapshots of the milling progress are shown in Figure 4d. These images are captured upon interupting the FIB milling, with the foil side of the lamella in focus. The distance *w* changes as the lamella thins, and the extracted instantaneous thickness *d* is indicated at the top of the snapshots along with the corresponding milling duration in color. In addition to these, we acquire a series of RLM images *during* FIB milling, with the optical focus set halfway between bottom and top sides of the lamella. We extract the mean reflected intensity along the *Y* direction as shown in Figure 4e, for different milling time points. Two characteristic intensity peaks at the start (*Y* = ∼7 µm) and the end (*Y* = ∼25 to ∼20 µm) of each line trace originate from respectively the bottom and top side of the lamella. By determining the width *w* in each line trace (or respective image), the lamella thickness can be calculated and plotted as a function of milling duration (Figure 4f). The insets show the first and final images from which *w* was determined. This illustrates how the milling geometry in the RLM images allows monitoring of the milling process down to a thickness of about 1 µm.

Thin-film interference effects can be exploited to determine the lamella thickness more accurately than with geometric considerations, as pioneered by Last *et al.*, who used multicolor RLM to determine the thickness at the periphery of vitrified cells grown on cryo-EM support films [20]. We illustrate the use of thin-film interference for thickness determination in our coincident light microscope by first fabricating a rough milled lamella of approximately 2.6 µm thickness, as shown in Figure 5a. We then defined a cleaning cross-section pattern in the FIB field of view in order to thin the lamella at a constant rate (Figure 5b, top). This pattern was milled line by line, with an effective milling rate of 2.4 nm*/*s along the FIB *Y* direction to ensure that the lamella is thinned evenly starting at the top (*t* = 0 s), and leaving a thin lamella after completion (Figure 5b, bottom and Figure 5c). The remaining thickness was approximately 430 nm, as determined from the width *w* measured in the RLM image. A discontinuous Pt GIS layer prevented further thinning beyond this point, illustrating the usefulness of Pt coating integrity read-out. Three different wavelengths were used for RLM image acquisition during FIB milling (*λ* = 463, 542 and 632 nm, 189 images per channel, 567 images in total), as shown in Figure 5d. For each wavelength, 35 individual line traces were extracted from the white rectangles, and plotted against the lamella thickness. The reduction in lamella thickness was computed from the image timestamp and the constant milling rate of 2.4 nm*/*s. A lamella thickness offset of 390 nm was used when plotting the traces, which is within the error of the estimate based on geometry, especially given that the protective platinum GIS layer is non-uniform. The dashed black line shows the reflectivity model *r*(*d, λ, T*), similar to [20]:

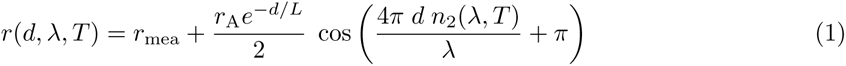

**Figure 5:**
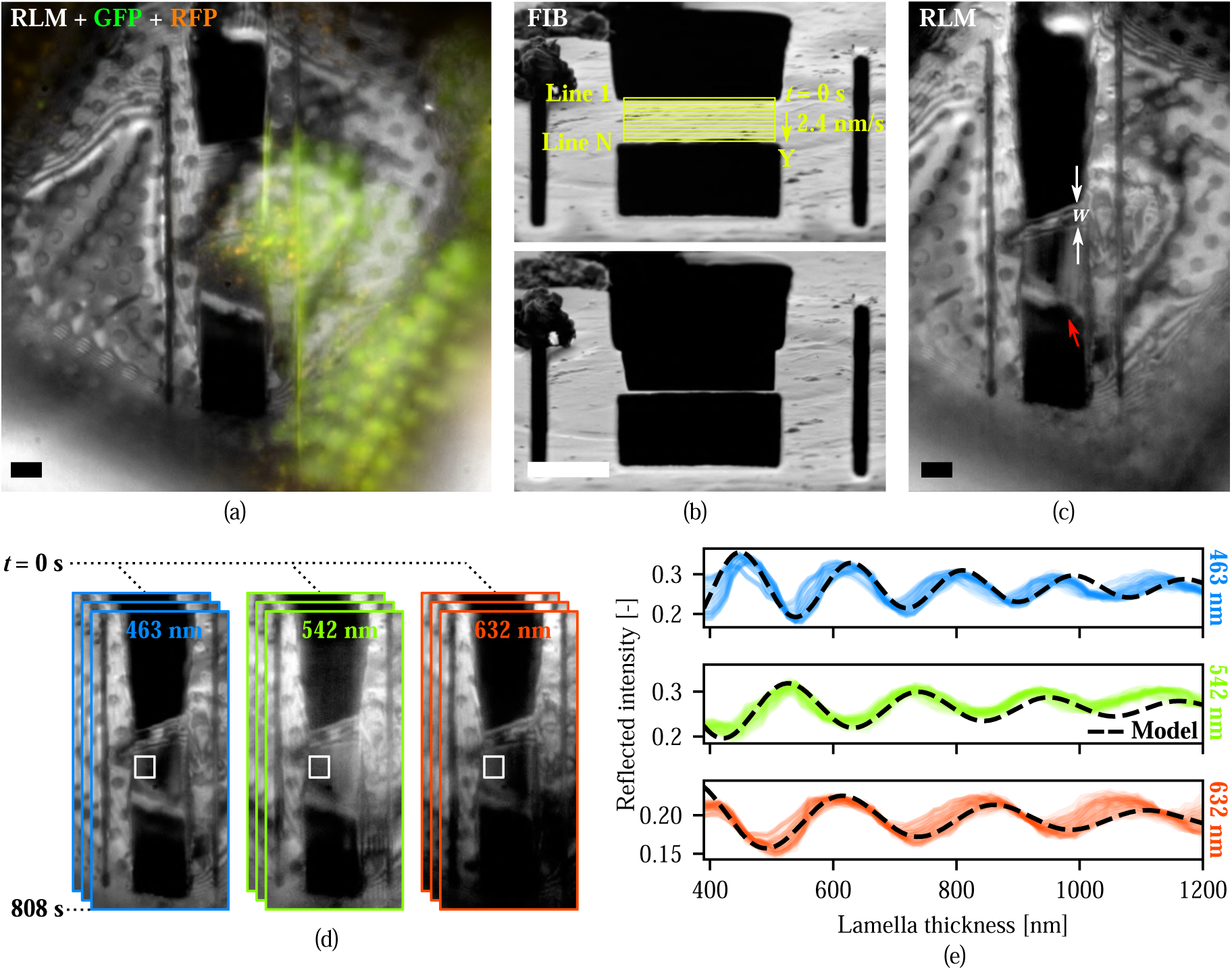
Reflected light intensity monitors lamella thickness during milling. **(a)** Lamella after rough milling, as imaged with fluorescence microscopy (colors) overlaid on reflected light (grayscale). At this stage, the lamella thickness *d* is approximately 2.6 µm (*w* = 15.3 µm). **(b)** The lamella after rough milling (top) and after thinning (bottom), as imaged with the FIB. The lamella is evenly thinned by defining a cleaning cross-section pattern in the FIB field of view (yellow marked area), having a constant milling rate of 2.4 nm*/*s along the FIB *Y* direction. The FIB milling progresses line-by-line at this rate, starting at the top (line 1) and ending at the bottom (line N)**(c)** Lamella after thinning, as imaged with reflected light (grayscale). At this stage, the lamella thickness *d* is approximately 430 nm (*w* = 2.5 µm), and the Pt GIS layer has been partially removed (red arrow), preventing further thinning. **(d)** Reflected light images are recorded in three different channels during the even lamella thinning. **(e)** For each reflected light channel, the normalized reflected intensity is plotted against the lamella thickness. 35 individual intensity traces are plotted for each channel (from white rectangle in (d)), along with the reflectivity model based on thin film interference (dashed black, Equation 1). All scale bars: 5 µm.

where *r*_mea_ and *r*_A_ are constants used to match the measured reflected intensity as determined from the acquired images. *d* is the lamella thickness, and *L* is a constant to account for scattering and loss of coherence occurring in the thin film, set to 500 nm. Corresponding values for the refractive index of amorphous ice *n*_2_(*λ, T*) are used from Kofman *et al.* [18].

Good agreement was obtained between the reflectivity model and the measured data for all three wavelengths, allowing the lamella thickness to be determined following Last *et al.* [20]. To reduce complexity, both in acquiring the measurement data and during analysis, we chose to measure the reflected lamella intensity for one wavelength only. By performing an error minimization on the reflectivity data, with the geometrical thickness estimate as an initial estimate, it is possible to generate 2D lamella thickness maps as a function of milling progression, as shown in the next section.

### Controlled lamella fabrication workflow

Finally, we combine the above in a single workflow for quality- and thickness-controlled fabrication of fluorescence-targetted lamellae. Initial target selection is done with fluorescence microscopy and quality control using the RLM, as discussed earlier. In reflected light imaging, the wedge-shaped endpoints of the lamella act as a Newton interferometer (NI) due to the changing thickness (Figure 6a). From an approximate lamella thickness of 1.5 µm, a region of constant thickness starts to appear in the middle, where thin-film interference (TFI) occurs. The normalized reflected intensity is recorded at a single point (blue cross) and is plotted using the geometrical thickness estimate *d*_est_ in Figure 6b (blue scatter). The thickness can now be further refined by error minimization, using the normalized version of the reflectivity model (Equation 1), yielding the green scatter in Figure 6b, and the green curve in Figure 6c. It shows more intricate thickness differences compared to the initial (geometrical) thickness estimate, partly because the thickness measurements stem from different locations on the section, but also as the (local) lamella reflectivity is more sensitive to thickness variations.

**Figure 6:**
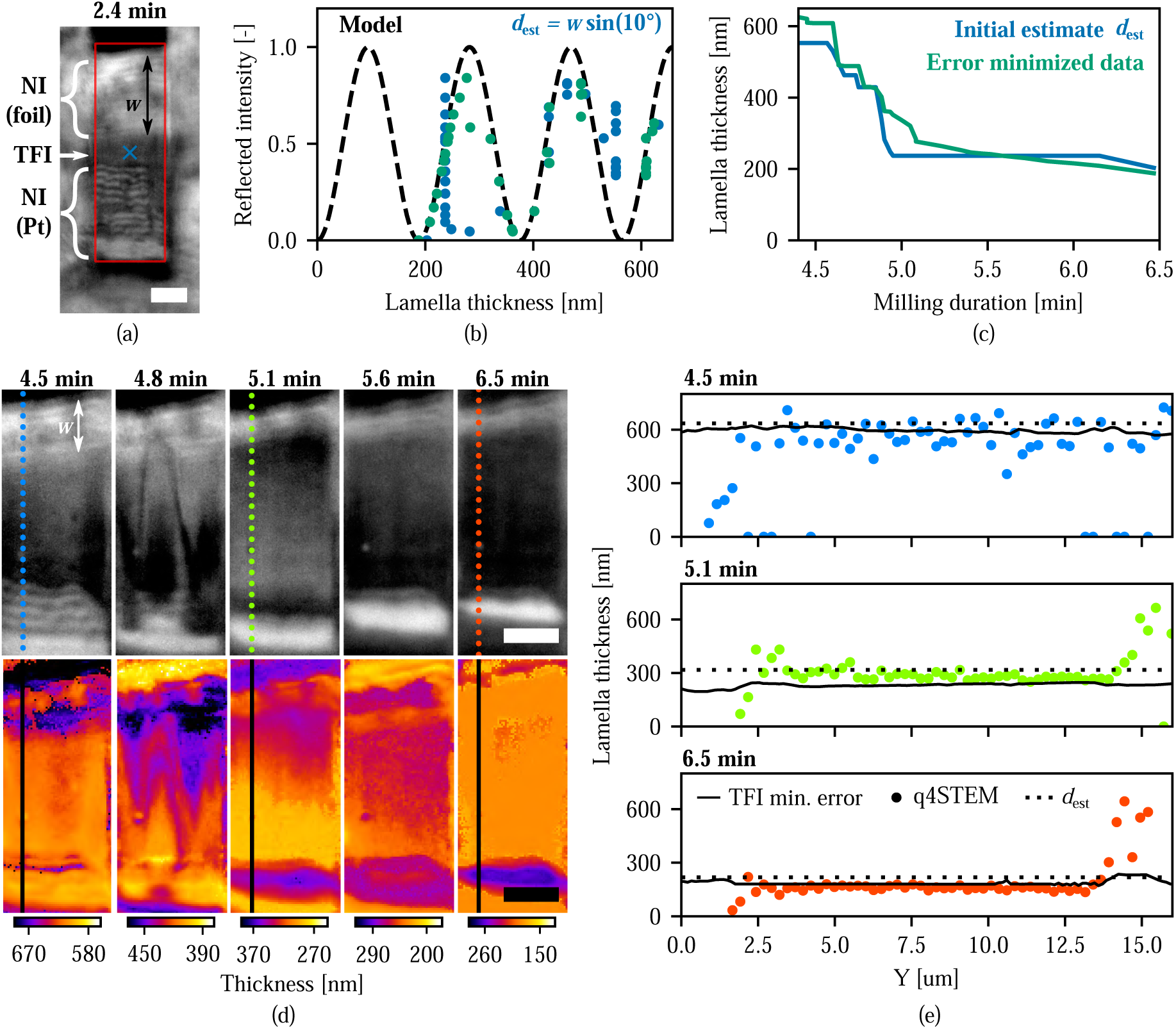
Thickness controlled lamella fabrication. **(a)** The lamella imaged in RLM (*λ* = 463 nm) at an intermediate milled state (*d ≈* 1.5 µm). The bottom (foil) and top (Pt GIS) sides of the lamella are wedge shaped and act as a Newton interferometer (NI). The region between has a more uniform thickness, and hence thin film interference (TFI) occurs. **(b)** With milling progression, the reflected intensity is recorded (at the blue cross in (a)), along with the lamella thickness estimate *d*_est_ = *w* sin(10*^◦^*) from the milling geometry (blue scatter). Error minimization is performed using a normalized version of the reflectivity model (dashed black), which outputs a refined lamella thickness estimate (green scatter), and finally yields **(c)** the lamella thickness versus milling duration. **(d)** The lamella imaged in RLM at varying stages of milling (top, grayscale). q4STEM data has been acquired at three time points during the milling (dotted colored lines). The error minimization process as outlined in (b) and (c) can be performed for all pixels in the RLM images, yielding thickness maps at varying stages of milling (bottom, colored). **(e)** Lamella thickness line traces along the *Y* direction, at three different time points. Thickness measurements include q4STEM (colored dots, see also (d) top), the error minimization of the normalized reflectivity (black lines, see also (d) bottom), and estimates from geometry *d*_est_ (dotted black lines). The increase in measured thickness around *Y* = 15 um is due to the increased density of the Pt GIS layer. All scale bars: 5 µm.

As reflected light *images* are acquired, this process can be extended for all the pixels on the lamella showing TFI, thus converting a gray scale reflected light image into a 2D thickness map of the lamella, as is done in Figure 6d. To visualize thickness variations at each stage of thinning the lamella, the minimum and maximum values of the color scale are changed for each image. At three time points during lamella thinning (4.5, 5.1 and 6.5 min), q4STEM data along the *Y* direction were acquired (colored dots). Along with the line traces of the 2D thickness maps (black lines), and the thickness estimated from the milling geometry (dotted line), the lamella thickness is plotted along the *Y* direction in Figure 6e. The three different methods are in overall good agreement, showing an approximate lamella thickness of 600, 300 and 200 nm at different stages of milling, but *d*_est_ overestimates the thickness below 300 nm. The mean difference between q4STEM and TFI measured thickness is *−*57, 56 and *−*21 nm for the three different time points, leading to a 10 to 20 % relative difference. At the 6.5 min time point a lamella thickness of 188 nm was reached, with a uniformity of *<*5 nm one sigma (from the error minimized TFI data). With the protective Pt GIS layer too thin and non-uniform, polishing was stopped.

The polished lamella was unloaded from the FIB-SEM using a cryogenic transfer system and loaded in a JEOL JEM3200-FSC TEM operated at 300 keV. An overview image (TEM settings) was acquired in the TEM, see Figure 7a. Fluorescence data, acquired after the polishing was finished, is overlaid on the overview image, and the alignment between TEM and the (reflected) light microscopy data is done in eC-CLEM [26]. This overlay is used to set the ROI for tomogram acquisition (Figure 7b, red box). A snapshot from the acquired tomogram (tomogram settings) is shown in Figure 7c (left), along with the segmentation overlay. The lamella thickness is measured from the *Y Z* slice along the dashed white line (c, right), yielding 189 nm, which agrees well with the 188 nm estimate from TFI.

**Figure 7:**
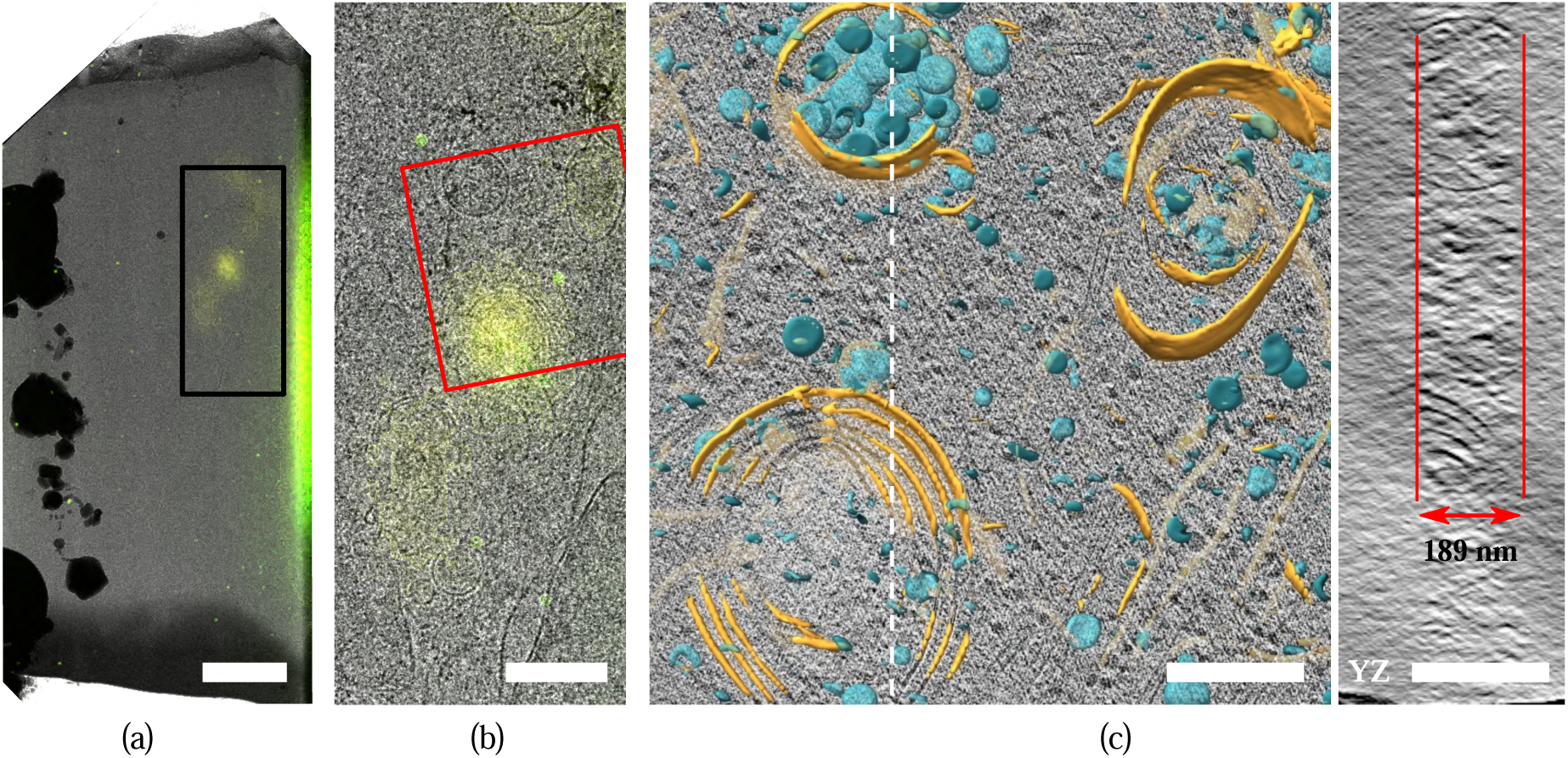
Cryo-electron tomography with fluorescence-targeted, thickness and quality controlled FIB-milled lamella. **(a)** Overview image of the lamella in TEM with overlaid fluorescence data. TEM and fluorescence images are aligned based on the reflected light image. This overlay is used for tomography target selection. **(b)** Zoom of the black-boxed area in (a). **(c)** Snapshot from the reconstructed tomogram with membrane segmentation as an overlay, acquired at the red rectangular area marked in (b). A *Y Z* slice taken along the dashed white line is shown on the right, annotating the lamella thickness as measured from the reconstruction. Scale bars: **(a)** 2.5 µm, **(b)** 1.0 µm, **(c)** 250 nm.

## Discussion

We have presented three different methods to accurately determine the lamella thickness during FIB milling, using a light microscope integrated in a FIB-SEM system; (i) by selecting a scintillator better suited for use in a fluorescence microscope, we were able to apply an earlier presented quantitative 4D-STEM technique [31] to the cryogenic lamella fabrication workflow. Benchmarking this technique to EFTEM showed good agreement in the determined thickness. (ii) The lamella thickness was estimated during milling via the known milling and imaging geometry, which produced reliable estimates between 2000 to 400 nm. (iii) Exploiting thin-film interference phenomena on a per-pixel basis yields a fine-grained thickness map of the lamella, thus providing quantitative feedback on the lateral thickness variations. The final lamella thickness estimate was compared with the thickness determined from reconstructed TEM data and showed excellent agreement.

We further showed how to target fluorescent regions of interest whilst correcting for the axial distortion induced by the inherent presence of a refractive index mismatch when imaging into a frozen cell. RLM is used to inspect milling sites to discard those sites which have parasitic ice present underneath the EM grid. A rudimentary form of communication between the FIB and FM control PCs is used to perform automated rough milling of fluorescent targets.

For different sample types, such as specimen prepared by the Waffle method or high pressure-frozen (HPF) the thickness can extend up to ∼200 µm [24, 16]. As such, depth dependent axial scaling effects need to be considered, and quantitatively corrected for by accurately measuring the axial coordinate *z_N_* (distance between refractive index interface and emitter) as observed in the optical microscope.

Our methods provide live feedback on lamella thickness and uniformity, integrity of the Pt coating layer, and presence of the fluorescent target when preparing a lamella out of a frozen hydrated cell. This reduces the complexity of the milling process and substantially increases the yield for producing lamella suitable for high-resolution cryo-EM analysis. Combined with the ability for reflection light and fluorescence-based targeting, our procedures pave the way towards fully automated and routine fabrication of frozen-hydrated cellular sections and highlight potential advantages of coincident FM-FIB-SEM instruments in workflow automation.

## 3 Limitations of Study

q4STEM and RLM thickness determination during the milling process require a coincident light microscopy setup integrated in the FIB-SEM system. This is absolutely required for the RLM measurements; for q4STEM an optional pixelated STEM detector may also be used and could provide improved sensitivity compared to the optical STEM signal detection employed in our setup. The q4STEM thickness determination method relies on Monte Carlo simulations of the scattering process of transmitted electrons and currently depends on parametric approximations of material composition and uniformity. Based on theoretical considerations these approximations may lead to thickness over- or underestimation of ∼ 2-3% for vitreous cellular material.

Alternative to a light microscope coincident with the FIB, an *in-situ* reflected light microscope might also be used for thickness determination through the geometry. Although this would require stage moves between the LM and acFIB, this can be implemented in an automated fashion in commercially available systems [32, 37].

## 4 Acknowledgments

We thank Ernest B. van der Wee for the useful discussions, and extend our gratitude towards Joyce van Loenhout and Leanid Kresik for providing cellular samples used during optimisation of automated FIB milling. We are thankful for the helpful discussions on the cryo-TEM workflow with the groups of C. Genoud and H. Stahlberg, during JPHs research stay at UNIL-EPFL. This work was financially supported by; NWO-TTW project N° 17152 to JPH.

## 5 Author Contributions

AJJ and JPH conceived and initialized the collaboration between their two research groups. First principle q4STEM experiments were carried out by RS, extended to the cryogenic workflow by DBB. DBB performed the experiments on thickness determination through the use of reflected light imaging. CT prepared the various cellular samples and together with DBB performed cryo-FIB milling. cryo-ET experiments were done by CT and WHE. DBB drafted the manuscript together with AJJ and JPH. All authors provided feedback and commented on the final version.

## 7 STAR Methods

### Resource availability

#### Lead contact

Further information and requests for resources should be directed to and will be fulfilled by the lead contact, Jacob Hoogenboom.

#### Materials availability

This study did not generate new unique reagents.

#### Data and code availability

All original code has been deposited at 4TU.ResearchData and is publicly available as of the date of publication. DOIs are listed in the key resources table.

### Method details

#### Cell culture

Briefly, cells were cultured in Roswell Park Memorial Institute 1640 Medium (RPMI, Gibco) supplemented with 10 % Fetal bovin serum (FBS) at 37 *^◦^*C with 5 % CO_2_. Glow-discharged carbon-coated gold mesh grids (QF 2/2 AU 200, Quantifoil) were placed on 3D printed holder (see [11]) and cells were gently dispensed onto it to reach 40 to 50 % cells confluency. Subsequently, cells were starved for 2 h using Hanks’ Balanced Salt Solution (HBSS, Gibco). For cryo-protection prior to plunge freezing we adapted a protocol by [15]: After removal of HBSS, cells were submerged in medium supplemented with cryo-protectant at increasign concentrations (RPMI/10 % FBS with DMSO/Glycerol 1.75 %, DMSO/Glycerol 2.5 %, DMSO/Glycerol 3.5 % and DMSO/Glycerol 7 %). Grids were removed from the holder, 2 µL of the final cryo-protectant solution was added on the back side before placing the grid on a Leica GP2 vitrification robot. Grids were blotted from the back for 12 s in the chamber maintained at 98 % humidity and finally plunged into liquid ethane. Grids were clipped in Autogrid cartridges with milling notch before transfer to the FIB/SEM holder.

#### Microcrystal formation

Hen egg-white lysozyme microcrystals were grown using room-temperature batch crystallization. Briefly, hen egg-white lysozyme was dissolved to 50 mg*/*mL in 20 mM sodium acetate pH 4.6. 100 µL of this solution were then mixed in a microcentrifuge tube with 250 to 400 µL of precipitant solution consisting of 18 % (w/v) NaCl, 6 % (w/v) PEG 6000, 0.1 M sodium acetate pH 3.0. Lysozyme microcrystals formed immediately as evidenced by the turbid solution. Samples using different protein to precipitant ratios were screened using negative staining TEM to optimize conditions for homogeneous microcrystal formation with dimensions smaller than 5 µm. A 1:4 (v/v) ratio of protein to precipitant solution was found to yield the most homogenous microcrystal populations. For vitrification, 3 µL of freshly prepared microcrystal slurry was applied on the carbon side of a glow-discharged QF R2/1 200 mesh grid (Quantifoil) kept at room temperature and 98 % humidity on a Leica GP2 vitrification robot, blotted from the carbon side for 12 seconds and flash-frozen in liquid ethane. Prior to loading in the FIB-SEM, the grids were mounted in FIB-compatible AutoGrid cartridges and secured with a c-clip (Thermo Fisher Scientific).

#### Lamella fabrication

Lamellae were fabricated using the focused ion beam on a Helios Nanolab 650 (Thermo Fisher Scientific) operating at 30 kV. Rough milling started at 2.5 nA and when decreasing lamella thickness the current decreased to 0.79, 0.23 and 0.08 nA.

#### Energy-filtered TEM

Thickness determination in EFTEM was done on a JEOL JEM3200-FSC operated at 300 kV by acquiring unfiltered and zero-loss filtered images of the lamella at 2500 *×* magnification (3.59 nm per pixel) on a Gatan K2 Summit direct electron detector either with- or without a 20 eV slit inserted below the omega filter. Montages of the lamella were generated with PySerialEM [33]. Aligned montages were used to compute *I/I*_0_ by dividing the pixel values of the zero-loss image by those of the unfiltered image. A thickness map was finally obtained by mapping 320 nm *·* ln(*I*_0_*/I*) onto the lamella montage, where a literature value of 320 nm for the inelastic mean free path of 300 kV electrons in vitreous ice was used [44].

#### Cryo-electron tomography

Lamellae were prepared as described above and stored in liquid nitrogen prior to imaging. Tilt series was acquired on a JEOL JEM3200-FSC operated at 300 kV with a bidirectional tilt scheme starting from 0 to *−*60*^◦^* and subsequently from 2 to 60*^◦^*, with a 2*^◦^* tilt increment. At a magnification of 6000 *×*, the pixel size was 6.3 Å and a defocus of *−*3 µm was used. The tilt series was aligned using patch tracking and reconstructed using weighted back-projection as implemented in AreTomo [45]. Segmentation was done in EMAN2 using a machine learning approach [5]. Correlation between LM images and micrographs was performed using the ecCLEM plugin in ICY software [25]; tomogram and volume visualisation was done in ChimeraX [12].

#### Scintillator selection for low temperatures

In Table S1, YAG is listed with other scintillator candidates are listed having a specified optical emission spectrum shifted towards lower wavelengths. LuAP was ordered (0.17 mm thick, 0.1 atomic percent cerium doped LuAlO_3_, single crystal, (110) oriented, Surface Preparation Laboratory B.V.) and several 3.4 mm diameter discs were laser cut. To avoid charging effects during electron beam exposure, the scintillator was coated with a 40 to 50 nm thick carbon film by means of thermal evaporation deposition. With an approximate specimen to scintillator distance of 300 µm, a maximum scattering angle of *θ_u_* = 218 mrad can be measured [31]. We have characterized how the electron induced scintillation yield changes as a function of temperature. A fiber coupled spectrograph (Princeton Instruments Acton SP2156 with a PyLoN:100BR Excelon camera) is connected on the excitation lights source (LS) input on the optics module, and the dichroic filter (DF) is replaced with a 50/50 beamsplitter (for details on the optical components see [3]). This allows collection of scintillator emission spectra whilst also imaging the scintillation spot on the camera. A primary electron beam energy of 5 keV is set along with a 0.1 nA beam current. Emission spectra are recorded whilst cooling down from 300 to 110 K as shown in Figure S1b, top. No spectral shift is recorded, but the maximum intensity decreases by a factor of 2.5 (Figure S1b, bottom). LuAP, given its orthorhombic crystal structure, introduces birefringence into the optical system, which is filtered out by adding a polarizing beamsplitter (PBS) in front of the camera (water-cooled Andor Zyla 4.2 PLUS). Depending on the orientation of the PBS either the parasitic pattern originating from the scintillator birefringence or the electron induced scintillation spot can be imaged, see Figure S1c. Without PBS present in the optical path, both are seen simultaneously (top).

#### Key Resources Table

| REAGENT or RESOURCE | SOURCE | IDENTIFIER |
| --- | --- | --- |
| Data and algorithms |  |  |
| All data and analysis scripts reported in this paper, are found at | 4TU.ResearchData | 10.4121/aeb7cf17-1b4a-4f09-a2fb-88d7875c672d |

## Supplemental information

**Table S.1:** Overview of scintillator candidates.

| Scint. | Cryst. struct. | $\lambda_{ab}$ [nm] | $\lambda_{em}^{max}$ [nm] | BF | Ref |
| --- | --- | --- | --- | --- | --- |
| YAG | C | 460, 340 | 550 | No | [8] |
| YAP | OR | 300 | 370 | Yes | [8, 9] |
| LYSO | MC | 250, 360 | 425 | Yes | [8, 2, 27] |
| LuAP | OR | 300 | 365 | Yes | [8, 10, 21, 34] |

C, OR and MC refer to cubic, orthorhombic or monoclinic, BF indicates birefringence.

**Figure S.1:**
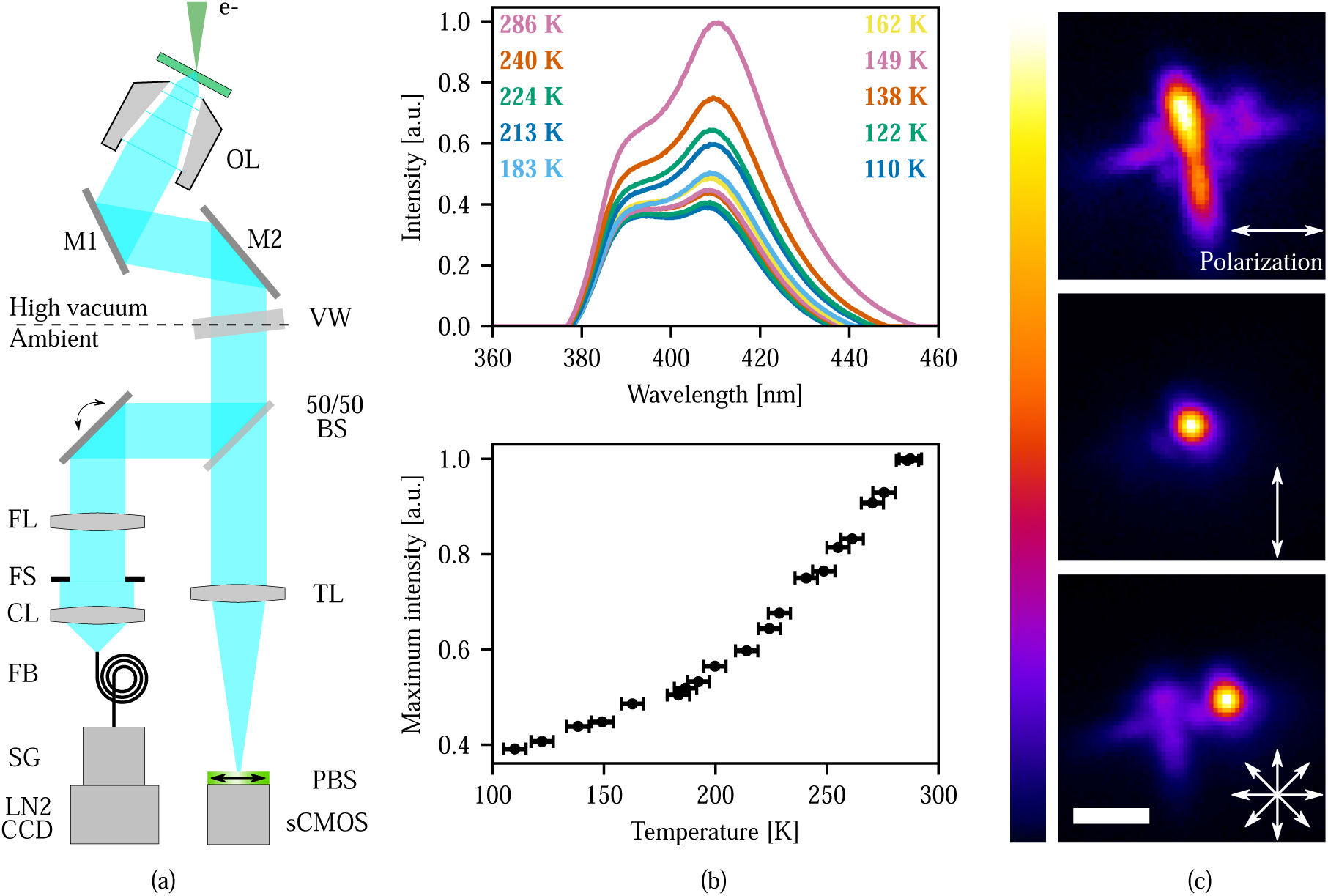
Optical properties of a LuAP scintillator under electron irradiation and the layout of the optical path. **(a)** Overview of the optical path denoting relevant components. The LuAP scintillator is irradiated with 5 keV, 100 pA electons. The scintillation light is collected by the objective lens (OL) and passed to the 50/50 beam splitter (BS), where it is divided between the excitation- and emission arms. This allows part of the light to be passed to the tube lens (TL) and focussed onto the camera (sCMOS) to form an image. A polarizing beam splitter (PBS) is added in front of the camera. The fiber coupled light source is replaced with a spectograph (SG) fitted with a liquid nitrogen cooled CCD camera. **(b)** Electron induced scintillaton emission spectra as a function of scintillator temperature (top). Peak positions do not shift, but the scintillation yield decreases. Peak maxima are plotted against temperature (bottom). The scintillation yield decreases with a factor of 2.5 going from 286 to 110 K. Horizontal errorbars of 5 K. **(c)** Scintillator birefringence is visualized by imaging the electron induced scintillation spot with the optical system. Without PBS present two spots are present (top) and the parasitic spot (middle) can be filtered out (bottom) depending on the PBS orientation. Scale bars: **(c)** 2 µm.

**Figure S.2:**
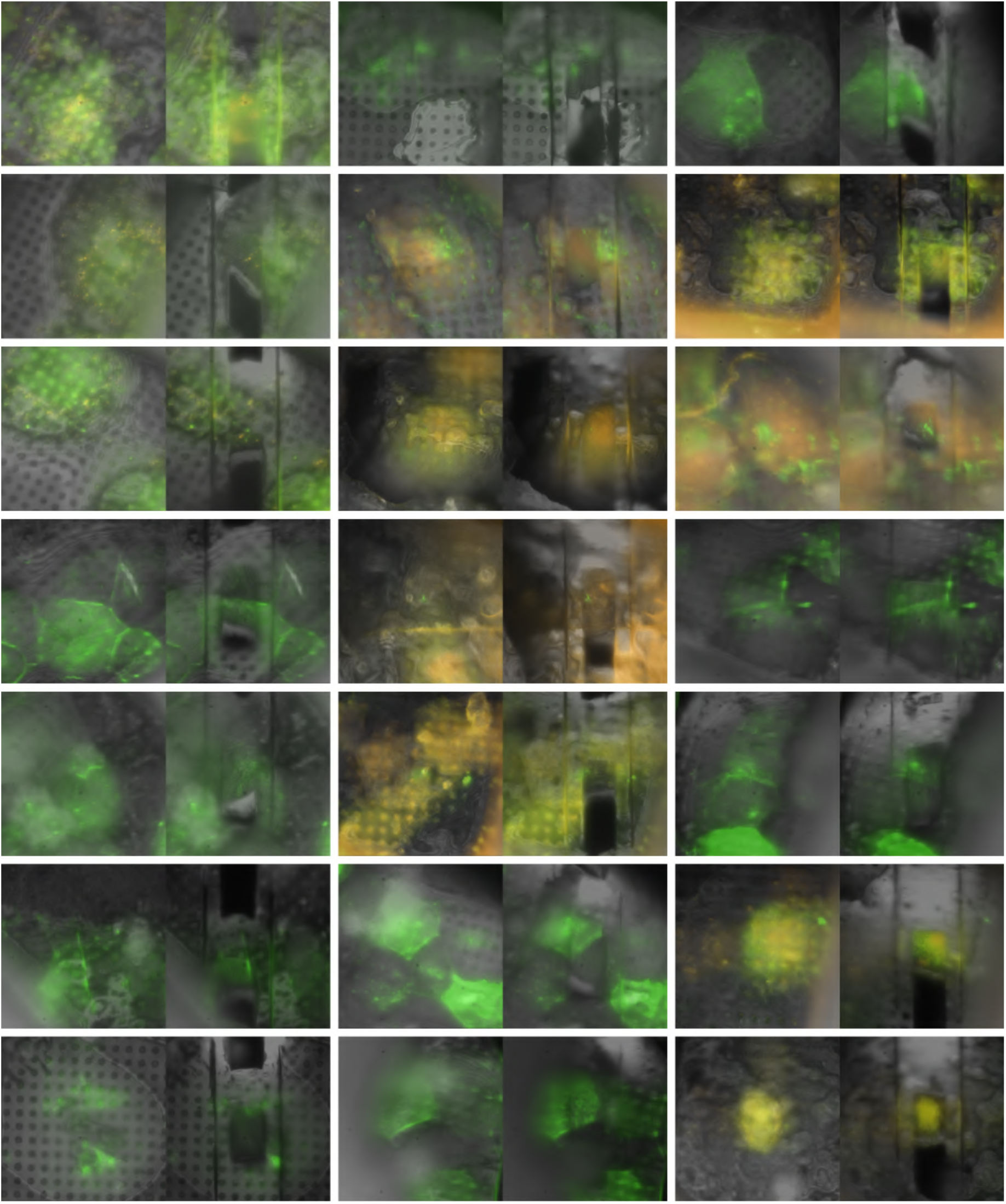
Fluorescent (colors) overlaid on reflected light (grayscale) image pairs. For each set, the left image is acquired prior to rough milling and the right image after. Scale bar 10 µm.

**Figure S.3:**
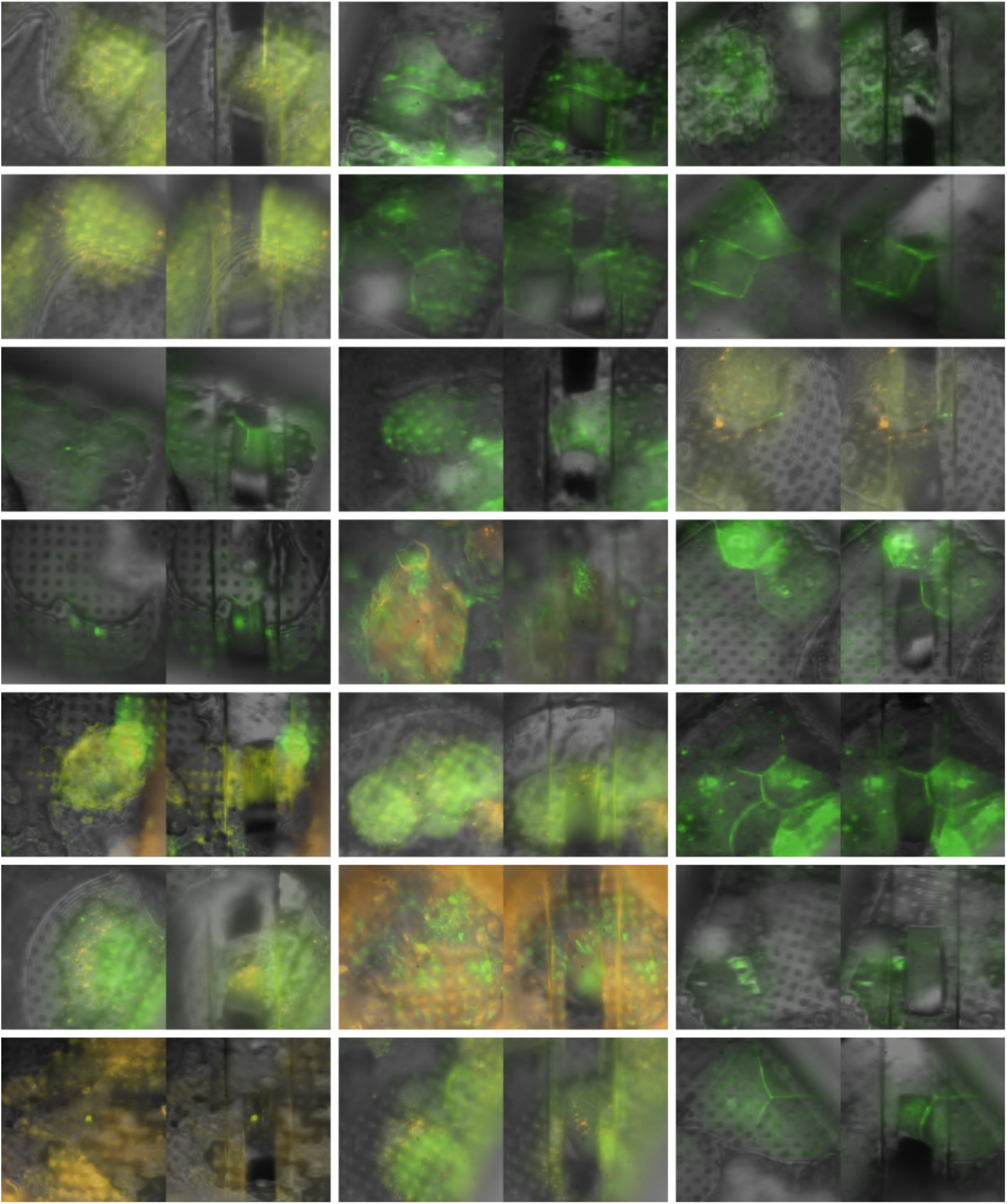
Fluorescent (colors) overlaid on reflected light (grayscale) image pairs. For each set, the left image is acquired prior to rough milling and the right image after. Scale bar 10 µm.

**Figure S.4:**
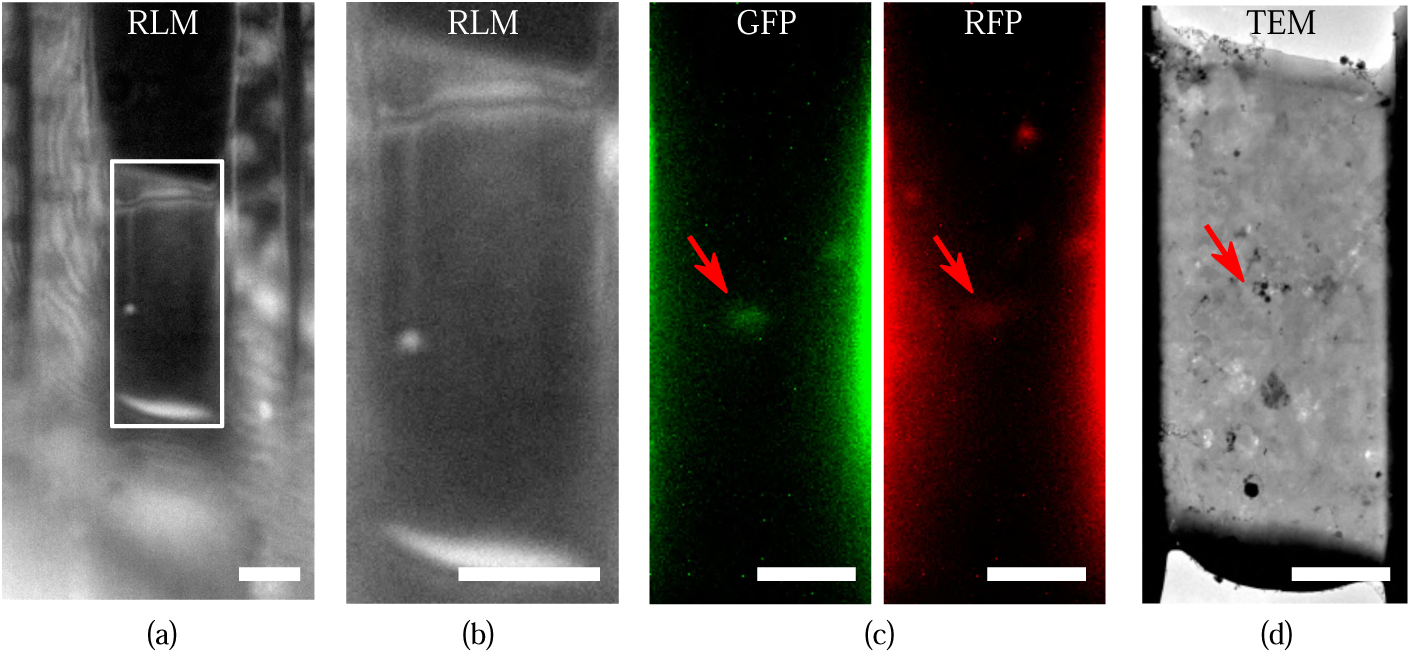
Fluorescence targeted lamella as fabricated for q4STEM thickness determination and verification with EFTEM. A red fluorescent protein (RFP)-GFP tandem fluorescent-tagged LC3 (mRFP-GFP-LC3) single molecule-based probe is used that can monitor the autophagosome maturation process [17]. **(a)** lamella as imaged with RLM after thinning with the FIB, and in **(b)** zoom from white rectangle. **(c)** Separated fluorescence microscopy channels, imaged after polishing (white rectangle from **(b)**), with target present (red arrows). From left to right: GFP emission (525 nm in green), and RFP emission (607 nm in orange). **(d)** Overview image as acquired in the TEM. All scale bars: 5 µm.

**Figure S.5:**
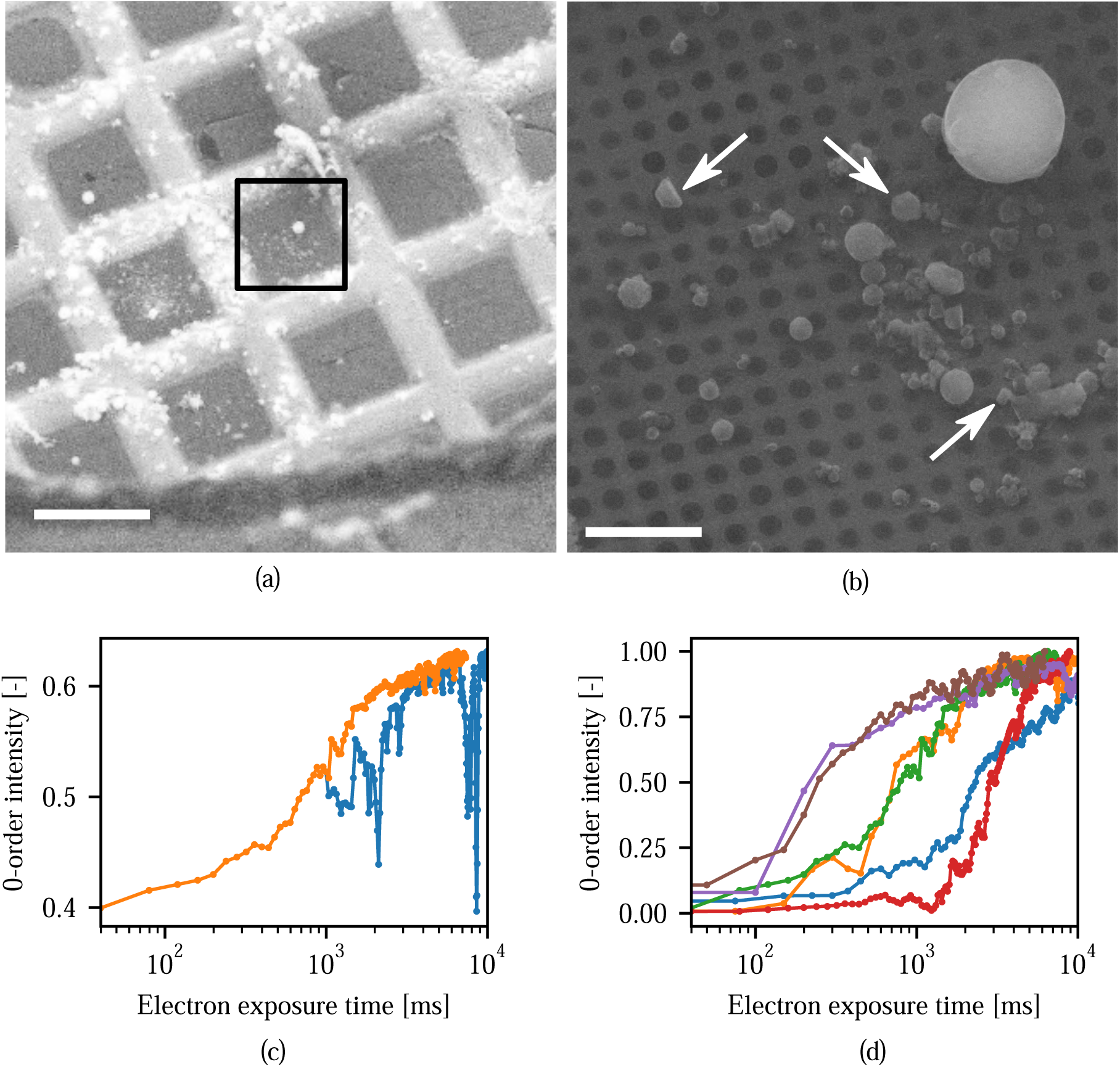
Assessment of radiolytic damage by q4STEM measurements. **(a)** Overview image of the EM grid containing small protein crystals. **(b)** The area as indicated in (a, black rectangle) shown at a larger magnification. White arrows denote the crystals which showed (partial) diffraction patterns. **(c)** With the optical focus set to the scintillator, the electron beam was moved over each crystal in search of a (partial) diffraction pattern. Once found, a time-lapse acquisition was started, from which the 0-order intensity is plotted against electron exposure time. The discontinuities seen in the blue trace originate from ∼ 2 to 4 nm peak-to-peak sample vibrations, causing slightly different parts of the crystal to be exposed [3]. By enforcing a monotonic increase, we filter and remove these discontinuities (orange trace). **(d)** Filtered 0-order intensity peaks for different crystals, as indicated by the white arrows in (b). Scale bars: **(a)** 100 µm, **(b)** 10 µm.

